# An integrative approach unveils a distal encounter site for rPTPε and phospho-Src complex formation

**DOI:** 10.1101/2021.01.13.426511

**Authors:** Nadendla EswarKumar, Cheng-Han Yang, Sunilkumar Tewary, Yi-Qi Yeh, Hsiao-Ching Yang, Meng-Chiao Ho

**Author notes:** Corresponding Author’s e-mail address, Meng-Chiao Ho, Hsiao-Ching Yang.

## Abstract

Protein tyrosine phosphatase: phospho-protein complex structure determination, which requires to understand how specificity is achieved at the protein level remains a significant challenge for protein crystallography and cryoEM due to the transient nature of binding interactions. Using rPTPεD1 and phospho-SrcKD as a model system, we established an integrative workflow involving protein crystallography, SAXS and pTyr-tailored MD simulations to reveal the complex formed between rPTPεD1 and phospho-SrcKD, revealing transient protein–protein interactions distal to the active site. To support our finding, we determined the associate rate between rPTPεD1 and phospho-SrcKD and showed that a single mutation on rPTPεD1 disrupts this transient interaction, resulting in the reduction of association rate and activity. Our simulations suggest that rPTPεD1 employs a binding mechanism involving conformational change prior to the engagement of cSrcKD. This integrative approach is applicable to other PTP: phospho-protein complex determination and is a general approach for elucidating transient protein surface interactions.

## Introduction

Protein-tyrosine phosphorylation is a reversible post-translational modification that regulates cellular signaling in eukaryotes. Protein-tyrosine phosphorylation levels in the cell are balanced by counteracting activities between protein-tyrosine kinases (PTKs) and protein-tyrosine phosphatases (PTPs)^1^. Aberrations in the regulation of protein-tyrosine phosphorylation are often associated with disease states such as arthritis, diabetes and cancer^1–6^. Crystallographic and peptide-binding studies of various PTPs such as PTP1B, SHP-1, SHP-2, rPTPε and rPTPα have revealed detailed mechanisms of substrate specificity/recognition at the active site^7–12^. The hallmark of previous structural studies is that the cysteine-dependent active site typically features a small, deep pocket to accommodate the phosphorylated tyrosine (pTyr) side chain and a relatively flat outer surface for the adjacent residues^13^. The interactions between the pTyr side chain and the active-site pocket provide most of the binding energy and drive the binding event. However, previous studies of the rPTPα phosphatase domain (rPTPαD1) and pTyr peptides with sequences derived from its physiological substrate, Src, displayed an unlikely weak affinity, with Michaelis constants (K_M_) in the millimolar range- much higher than the physiological concentration^10^. Although the D2 domain of rPTPα and SH2 domain of Src also play crucial roles in rPTPα: Src complex formation^9,14,15^, studies of ERK kinase and metalloproteinase have shown that additional protein-protein interaction (also known as encounter interface or exosites) far from the active site can facilitate substrate recognition^16^. Currently, there is only one PTP: phospho-protein complex structure in protein data bank (PBD), but it represents a noncatalytic mode of interactions and cannot reveal additional protein-protein interactons^17^. Therefore, the corresponding encounter interface in PTPs remained largely unexplored as no functional PTP: phospho-protein complex structure has yet been determined.

Herein, we report the first rPTPεD1: phospho-SrcKD complex structures by integrating experimental and computational approaches that is applicable to other PTP complexes. In brief, the experimental SAXS data guides rigid-body docking to form the initial complex, which provides a defined spatial orientation between rPTPεD1 and phospho-SrcKD. This approach effectively reduces the computational time and resource required by multiscale MD simulations in searching of protein-protein binding ensemble structures ^18–21^. The following pTyr-tailored MD simulation optimized the spatial arrangement of the two protein molecules and the encounter interface. The key residues and trajectory snapshots of protein complex formation are further revealed by steered MD simulations and umbrella sampling.

Our complex structure revealed an encounter interface, which greatly enhance the formation of a catalytically competent complex. A single site was replaced on the encounter interface, designed to partially disrupt charge-charge interactions, resulting in a seven-fold reduction of the association rate *k*_on_, and a 30% reduction of PTPε phosphatase activity towards phospho-SrcKD but not towards pNPP, a pTyr substrate analog. Our structural analyses further suggest that a conformational selection mechanism plays an initial role in molecular recognition between rPTPεD1 and SrcKD.

## Results

### Production of stable rPTPεD1: phospho-SrcKD complex

In the present study, we focused on the interaction between the D1 domain of rPTPε (rPTPεD1) which possesses phosphatase activity and its target, the kinase domain of Src (SrcKD) where the C-terminal pTyr527 is dephosphorylated. A known catalytically inactive and substrate-trapping mutation of rPTPεD1-C335S was used to obtain a stable rPTPεD1: phospho-SrcKD complex. CSK, a known kinase of the Src family, was used to phosphorylate SrcKD *in vitro*^22^. The SrcKD double variant, K295M/Y416F substituted in ATP binding and activation residues, respectively, was produced to prevent additional auto-phosphorylation on Src^23,24^. The CSK treated SrcKD was pooled with rPTPεD1 for complex formation and was further purified by size-exclusion chromatography (SEC). The purified rPTPεD1: phospho-SrcKD complex was co-eluted at a volume that was distinct from that of the uncomplexed rPTPεD1 and SrcKD (Fig. 1a and 1b). The ability to be co-eluted in the SEC suggests that the phospho-SrcKD forms a stable heterodimeric complex. Our analytical ultracentrifugation (AUC) result shows that both uncomplexed rPTPεD1 and SrcKD show a single peak with an S20 value of ~ 3 whereas rPTPεD1: phospho-SrcKD complex shows an additional peak with S20 value > 4, indicating stable heterodimeric complex formation and consistent with SEC experiment (Fig. 1c).

**Figure 1.**
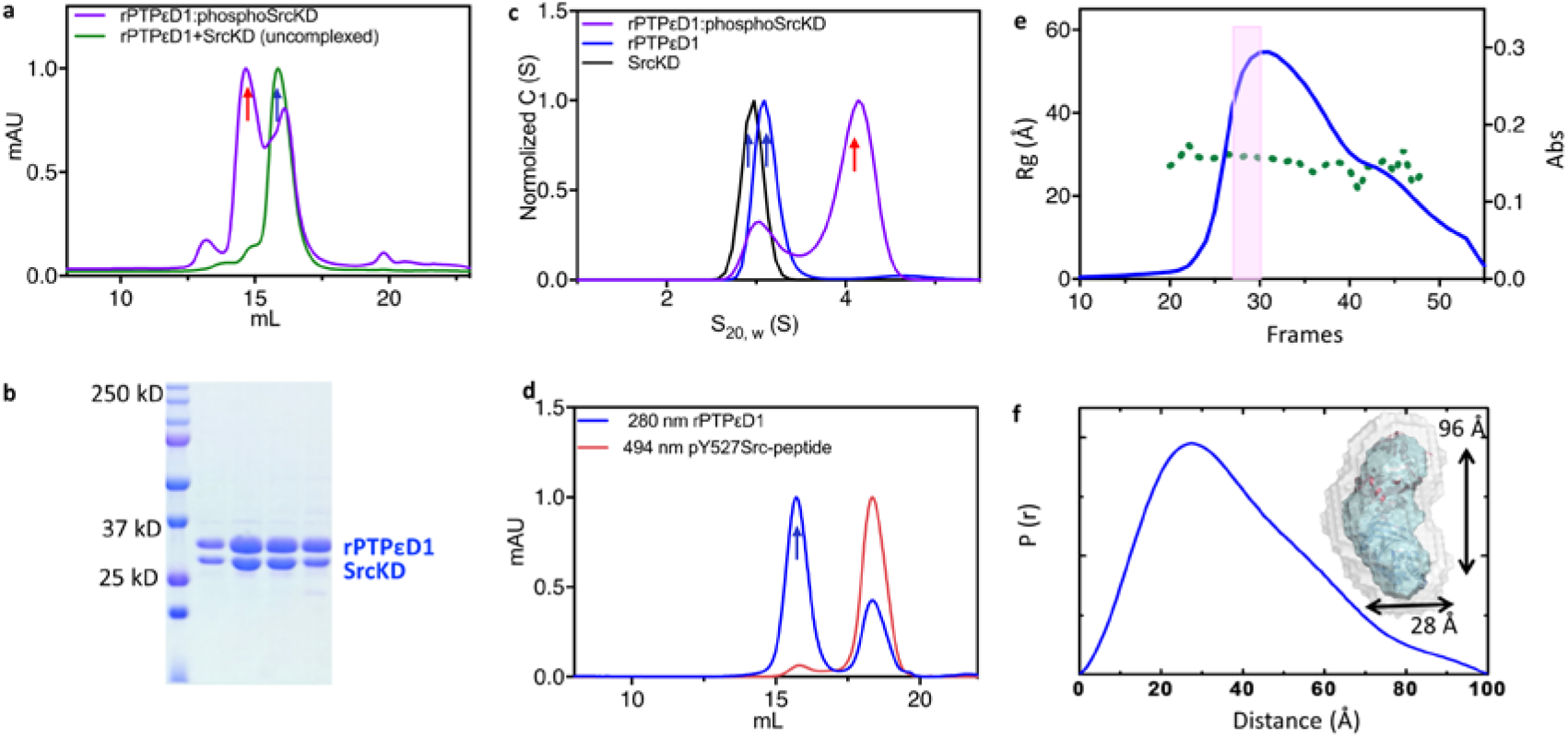
Characterization of the rPTPεD1: SrcKD complex. (a) The overlaid SEC profile of rPTPεD1: phospho-SrcKD complex colored in violet and a mixture of rPTPεD1 + unphosphorylated SrcKD (no complex form) colored in green. The elution position of complex and uncomplexed proteins are indicated as red and blue arrows, respectively (b) The SDS-PAGE result from SEC of rPTPεD1: phospho-SrcKD complex. The first lane is the reference marker with the corresponding MW shown. Lane 2-4 correspond to the highest peak area (red arrow) from SEC of PTPεD1: SrcKD complex shown in Fig. 2a. (c) The overlaid distribution of the sedimentation coefficient of rPTPεD1, phospho-SrcKD and complex are shown in blue, black and violet, respectively. The complex revealed an additional peak with S_20,w_ value of 4.1. The sedimentation coefficient of complex and uncomplexed protein are indicated as red and blue arrows, respectively. (d) Binding efficiency analysis of Src pTyr527 peptide towards rPTPεD1. The peptide is labeled with FiTC which can be detected by UV absorption at 484 nm wavelength (shown in red). The rPTPεD1 is eluted at 14 mL of elution volume as a peak with UV absorption at 280 nm (shown in blue and a blue arrow). The peptide is eluted at 18 mL elution volume and is not co-eluted with rPTPεD1. (e) R_g_ (green dots) extracted from the SEC-SAXS data measured along with the chromatogram (280 nm wavelength, blue line). (f) The pair-distance distributions P(r) computed from the scattering data from (a) plots of the rPTPεD1:SrcKD complex. Molecular envelope derived from SAXS data is shown.

### Distinct binding behavior of rPTPεD1 towards SrcKD and peptide

Previous findings revealed that rPTPα has a substantially weaker binding affinity (in the low mM range) toward the pTyr Src peptide^10^. As rPTPε is a homolog of rPTPα, SEC showing that rPTPεD1 does not co-elute with pTyr Src peptide is similarly a sign of weak or transient binding between rPTPεD1and Src peptide (Fig. 1d). As we observed stable rPTPεD1: phospho-SrcKD complex formation (Fig 1a), we hypothesized a non-peptide mediated binding regime and the existence of additional encounter interfaces (exosite) between rPTPεD1and SrcKD.

### Docking model by SAXS and MD simulation

The combination of multiscale MD simulations with solution SAXS is advantageous as MD simulations allow conformational arrangement while SAXS experiments provide information about overall shape which can effectively reduce the time-consuming simulation process in searching of protein-protein binding ensembles. SAXS (with a q value ranging from 0.009 to 0.2 Å^−1^) was used to determine a molecular envelope for the rPTPεD1: phospho-SrcKD complex, indicating an elongated particle in solution with a radius of gyration (Rg) of 29.4 Å and a maximum intramolecular distance, D_max_, of 91.1 Å (Fig. 1e, 1f, 2a and 2b and Table. S1). The calculated low-resolution envelope had adequate space to fit the complex molecules (Fig. 1f). In addition, the rigid-body docking complex was generated by CORAL using crystal structures of rPTPεD1, (PDB ID: 2JJD) and SrcKD (PDB ID: 2SRC). One of the best fit CORAL docking models with χ^2^ value of 4.1 showed a docking complex with tail-to-tail relative orientation (Fig. 2a). In this complex, there were no lysine or arginine residues found proximal to the encounter (intermolecular) interface. Further examination found that the complex cannot be cross-linked by amine-to-amine crosslinkers, such as glutaraldehyde and bis-sulfosuccinimidyl suberate (BS3), supporting this docking model.

**Figure 2.**
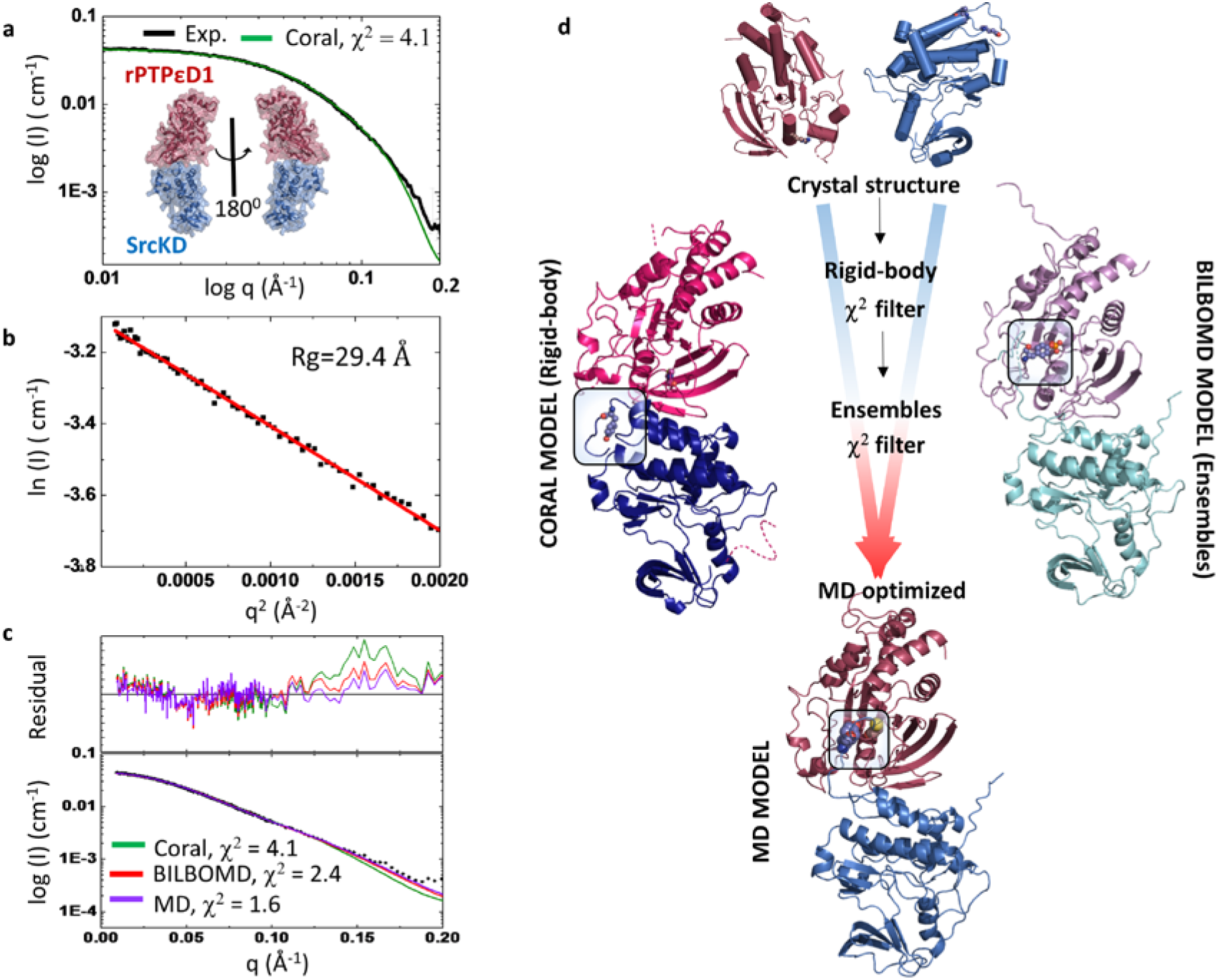
SEC-SAXS data and analysis of rPTPε:phospho-Src complex results. (a) (a) The simulated SAXS curve from the CORAL docking model (shown in green) is overlaid with experimental SAXS data on rPTPεD1: phospho-SrcKD complex (shown in black) with the χ^2^ of 4.1. The CORAL docking model of rPTPεD1: phospho-SrcKD. The rPTPεD1and SrcKD are colored in pink and blue, respectively. (b)The Gunier plot is shown with R_g_ of 29.4 Å. (c) Comparison of CORAL, BILBOMD and MD simulation model of rPTPεD1: phospho-SrcKD complex. The corresponding scattering profile and fitting of the experimental profile are overlaid. The χ^2^ calculated by FoXS is indicated. (d) The flow-chart of MD complex generation from the individual crystal structure. The individual crystal structure and CORAL, BIBLOMD and MD simulated complex models are shown. The rPTPεD1 and SrcKD are colored in magenta and blue system. The pTyr527 of SrcKD is highlighted in the box.

To create a pTyr bound docking model, we manually moved the flexible pTyr region (Asp518-Gln534) toward rPTPεD1 based on geometry restraints and positioned pTyr527 into the active-site of rPTPεD1 based on the pTyr-peptide bound PTP1B crystal structure (PDB ID: 1G1H)^25^. The ability of the C-terminal pTyr527 to reach the rPTPεD1 active site by moving only the flexible C-terminus implies that our tail-to-tail docking model is in a functionally competent state, allowing rPTPεD1 to dephosphorylate pTyr527 of SrcKD. The missing N-terminal residues (four residues in rPTPεD1 and 27 residues in SrcKD) were added to the CORAL docking model by RosettaCommons. By keeping pTyr527 bound in the active site and the N-terminus flexible, the docking model was optimized by BIBLOMD with an improved χ^2^ value of 2.5 (Fig. 2d and S1). However, close inspection of the BIBLOMD model revealed that Glu486 and Glu489 of SrcKD were surrounded by a negatively charged surface on rPTPεD1, indicating an unfavorable repulsive contact in the encounter interface (Fig. S2). Application of MD relaxation allowed these unfavorable repulsive contacts to be resolved into favorable attractive interactions in the encounter interface (Fig. 2d, S1 and S2). Compared to the crystal structure of uncomplexed rPTPεD1, the most apparent difference occurs in the N-terminal helices (residues 121-128 and 136-141) of rPTPεD1 that are more extended and undergo minor loop-helix-loop rearrangement. Those regions are distal to the encounter interface and there are no conformational changes of helical backbone observed in the vicinity of the complex interface of rPTPεD1. The side chain Arg220 present in the encounter interface rotates to the more solvent exposed side, providing an attractive favorable contact in the interface (Fig S1). In the case of SrcKD, a minor rearrangement of the backbone of one helix (residue 469-477) is observed. The major change is that the loop including Glu486 flips a distance of 4.5 Å toward the encounter interface, contributing to a favorable attraction in the encounter complex interface. Overall, the MD relaxation complex forms additional rPTPεD1-R220: SrcKD-E486 and rPTPεD1-K237: SrcKD-D518 interactions with χ^2^ values improved from 2.5 to 1.6, indicating a better fit to the experimental SAXS data (Fig. 2c, and Table S2).

### Mapping the interactions during complex formation with a free-energy approach

Typically, searching protein dissociation or association pathway requires long-timescale MD simulations combined with an additional modeling approach^26^, however, it is not easy to reach its convergence criterion. In contrast, our SAXS-based complex structure can quickly provids a reasonable initial complex model for further MD optimization.

Initially, MD simulation with umbrella sampling failed to assess the pathway trajectory owing to the strong attractive interactions between pTyr527 of SrcKD and the rPTPεD1 active site. Consequently, the complex remained in the bound form and resulted in significant rotation of the protein molecules (Fig. S3). Hence, the unphosphorylated form of SrcKD was purposely used for the following MD simulation.

The simulated reaction coordinate was selected based on the center-of-mass (COM) between rPTPεD1 and SrcKD. In the dissociation process, the encounter interface starts to disrupt at a COM distance of 49 Å and vanish at 55 Å (Fig. 3a). It suggests that the interface interaction of the complex started dissociating at COM distance of 49 Å and completely dissociated at 55 Å. To understand and evaluate the contribution of key residues, we decomposed the free energy of two charge-charge residue pairs, R220: E486 and K237: D518. The energy decomposition results suggested both residue pairs play roles in binding, which is consistent with the optimized MD model (Fig. 3b). In addition, our results illustrate that the rPTPεD1-R220: SrcKD-E486 pair show a larger difference between the bound state and the unbound state whereas the rPTPεD1-K237: SrcKD-D518 pair displays a minor change, indicating that the R220:E486 interaction plays a crucial role in complex formation.

**Figure 3.**
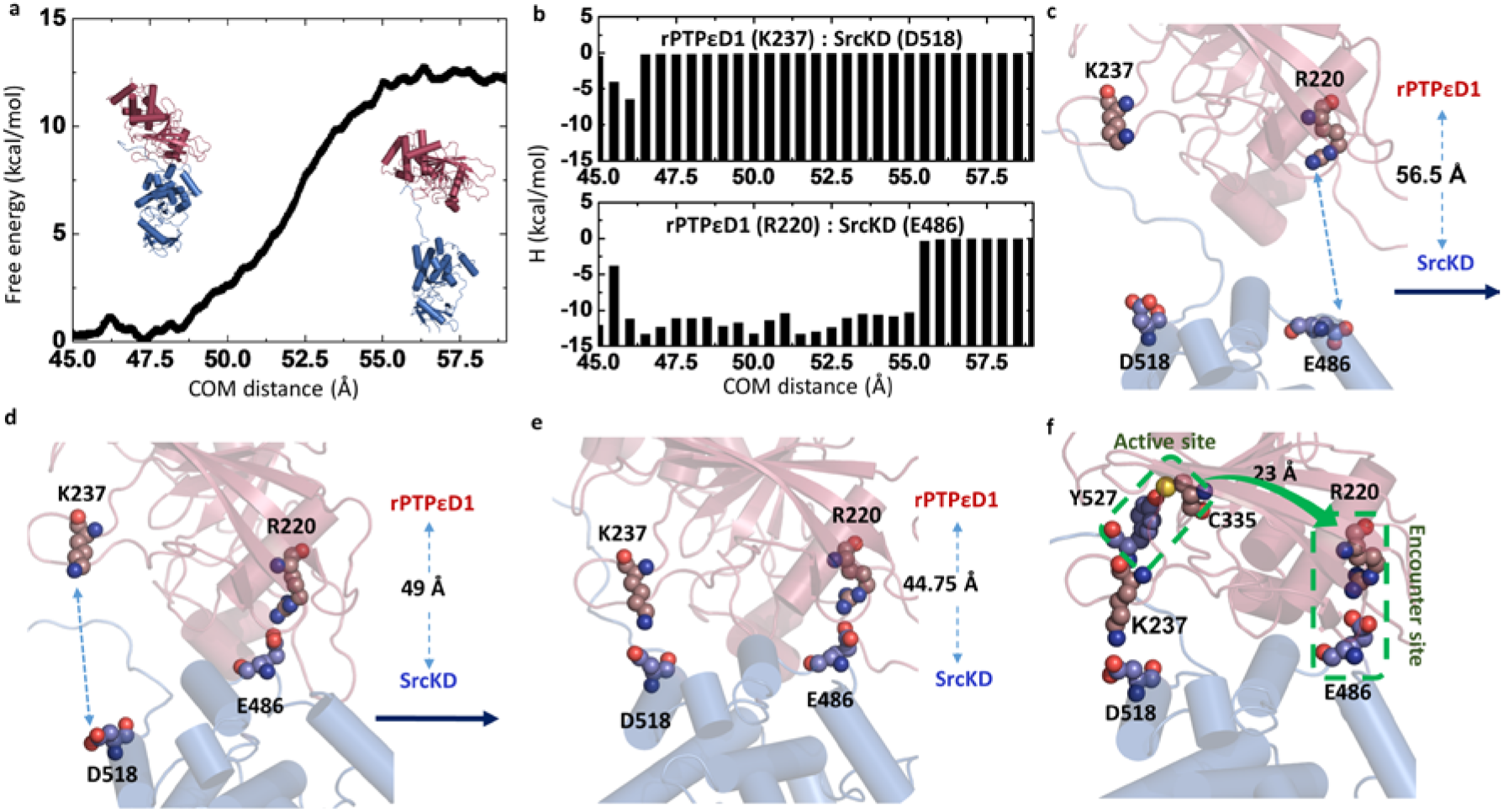
MD simulation reveals the trajectory snapshot along with the COM distance of the rPTPεD1:SrcKD complex formation. (a) In PMF approach, the free energy change along the COM distance revealed that unbound state in the COM distance was separated by 56.5 Å. (b) The energy decomposition by MM-GBSA was made to evaluate the contribution of three interacting residue pairs, The identified crucial interactions between the rPTPεD1:SrcKD complex is highly consistent with the MD optimized model. Also, the steered MD simulation mapping the interactions during complex formation. (c) In the unbound states, the complex initially formed a dynamic contact interface with the rPTPεD1 and SrcKD that COM distance is 56.5 Å. (d) Upon forming R220-E486 contacts, then K237-D518 contacts in the intermediate state. (e) Finally, the formation of these contacts facilitated the rearrangement of the complex to the bound state. (f) the proposed rPTPεD1: SrcKD docking model focused on the encounter site and active-site residues. The R220 is 23 Å away from the active-site residue, C335. The key interface residues and active-site residues are showed as spheres and labeled.

Next, to demonstrate the corresponding intermediate structural changes from the unbound to bound state, MD simulation in the association direction was performed. The starting model was derived from the previous MD pathway trajectory at the COM distance of 56.5 Å. To simulate the process of complex formation, proteins were slowly moved toward each other in the bulk solvent environment. In the unbound state, the proteins were very dynamic without any close contact (Fig. 3c). At the intermediate state, the complex gains electrostatic attraction between rPTPεD1-R220 and SrcKD-E486 (Fig. 3d). At the next stage of complex formation, an additional interaction is formed between rPTPεD1-K237 and SrcKD-D518 (Fig. 3e). SrcKD-D518 is only eight amino acid residues away from pTyr527 in the primary structure, so this intermolecular arrangement brings SrcKD-pTyr527 close to the rPTPεD1 active site (Fig. 3f). The identified interactions along the association pathway of rPTPεD1: SrcKD complex is highly consistent with the dissociation process results that R220:E486 and K237:D518 pairs play roles in the complex formation

### *In vitro* validation of the role of rPTPε-D1-R220 in complex formation

To further validate the role of rPTPεD1-R220: SrcKD-E486 interaction, a repulsive mutant rPTPεD1-R220E was generated. The SEC and AUC experiments showed that the rPTPεD1-R220E failed to form a stable complex with phospho-SrcKD (Fig. 4a and 4b). This result supports the model that the encounter interface of R220: E486 is crucial for the stabilization of the rPTPεD1-R220: phospho-SrcKD complex. The previous study of PTP1B shows disruption of charge-charge interactions within the pY-6 to pY+5 region of phospho-peptides decreases binding constants by 2 to 18 fold^27^. Our binding experiment showed that the R220E variant has a 7.8-fold reduction in association rate with phospho-SrcKD with *k*_on_ of 128 M^−1^s^−1^ and an estimated K_D_ of 257 μM compared to wild-type rPTPεD1 with *k*_on_ of 998 M^−1^s^−1^ and K_D_ of 34 μM (Fig. 4c and 4d). This relative slow association rate (well below the diffusion-controlled rate) indicates that the binding event is limited by conformational rearrangement^28^.

**Figure 4.**
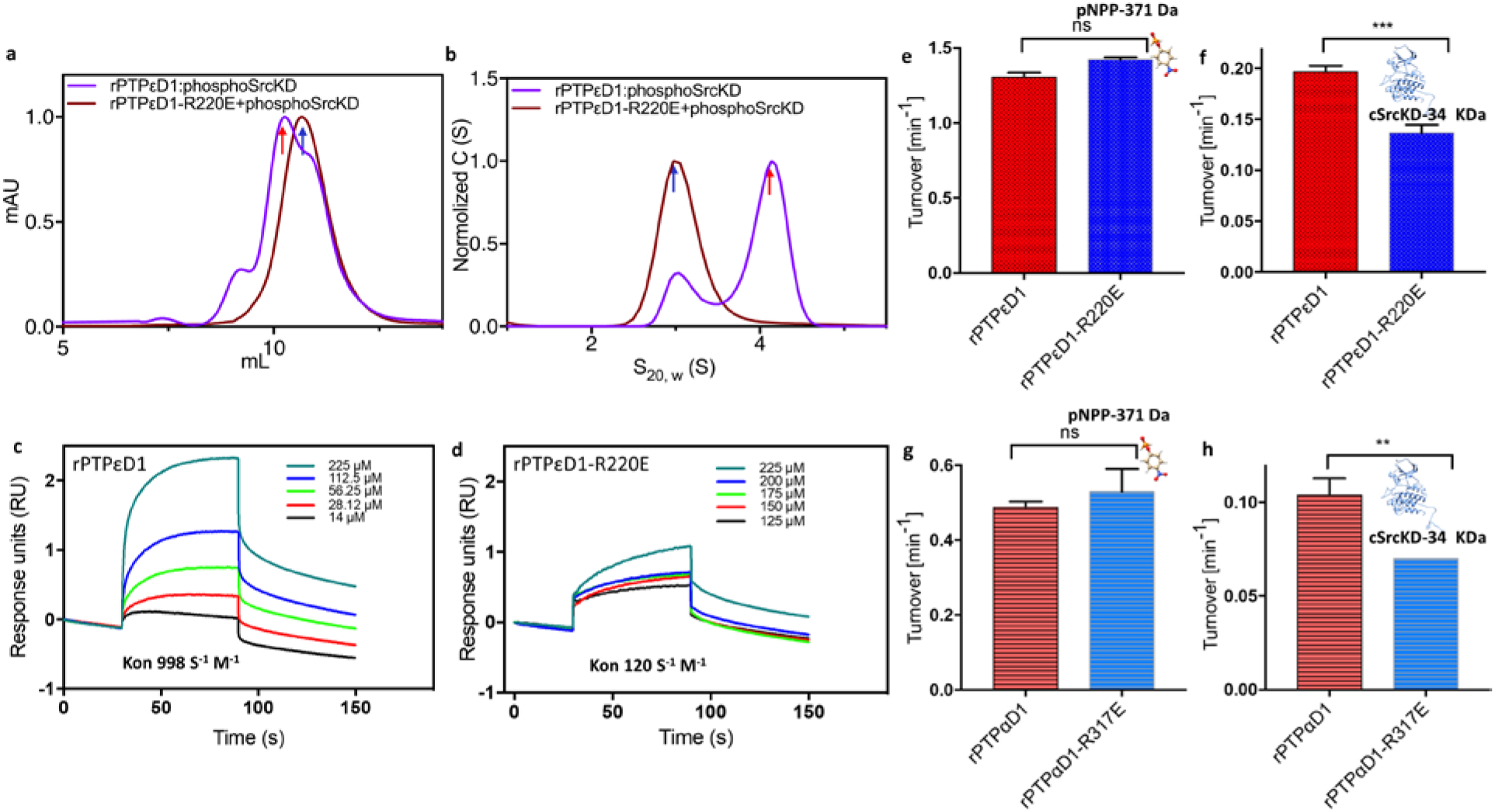
Comparison of rPTPεD1/rPTPεD1-R220E and wild-type toward phospho-SrcKD. (a) Elution profiles of rPTPεD1: SrcKD complex via SEC depicted in violet and of rPTPεD1-C335S/R220E:cSrcKD complex shown in brown. An excess amount of rPTPεD1 is added in rPTPεD1: SrcKD complex to show the elution volume of the uncomplexed form (the peak after 10 mL elution volume), which is overlaid with the peak of rPTPεD1-R220E: SrcKD complex, indicating transient complex formation. The elution position of complex and uncomplex are indicated as red and blue arrows, respectively. (b) The overlaid distribution of the sedimentation coefficient of rPTPεD1: SrcKD complex illustrated in violet and rPTPεD1-R220E: SrcKD complex is in brown. The sedimentation coefficient of complex and uncomplex proteins are indicated with red and blue arrows, respectively. (c) BLI sensogram of rPTPεD1 binding to the immobilized phospho-SrcKD protein. Various concentrations of rPTPεD1 are shown in different colors and labeled. The *k*_on_ value is indicated. (d) BLITz sensogram of rPTPεD1-R220E binding to the immobilized phospho-SrcKD protein. Various concentrations of rPTPεD1-R220E are shown in different colors and labeled. The *k*_on_ value is indicated. (e) The phosphatase activities of rPTPεD1 and rPTPεD1-R220E against pNPP is depicted in red and blue color, respectively. (f) Phosphatase activities of rPTPεD1 and rPTPεD1-R220E against phosphorylated SrcKD are illustrated in red and blue color, respectively. (g) The phosphatase activities of rPTPαD1 and rPTPαD1-R371E against pNPP is depicted in red and blue color, respectively. (h) Phosphatase activities of rPTPαD1 and rPTPαD1-R371E against phosphorylated SrcKD are illustrated in red and blue color, respectively.

Finally, we compared the phosphatase activity of the rPTPεD1 wild-type and R220E variant using pNPP and phospho-SrcKD as substrates. As expected, the pNPP assay results showed that both rPTPεD1 wild-type and R220E mutant possess similar phosphatase activity (Fig. 4e), suggesting the R220E mutation does not affect the catalytic site. A previous study of PTP1B activity toward phospho-peptide showed disruption of charge-charge interaction has little effect on *k*_cat_^27^. However, our phosphatase activity toward phospho-SrcKD revealed a ~30% activity reduction as measured by *k*_app_ for the rPTPεD1-R220E variant (Fig. 4f). Furthermore, sequence alignment shows that R220 is highly conserved in rPTPεD1 as well as rPTPαD1 (Fig 5a and 5 b). The rPTPαD1-R317E mutation (corresponding to R220E in rPTPε) also exhibited a ~30% decrease in activity compared to wild type rPTPαD1 (Fig. 4g and 4h), suggesting our proposed encounter interface may be conserved in the type IV subfamily of rPTPs.

**Figure 5.**
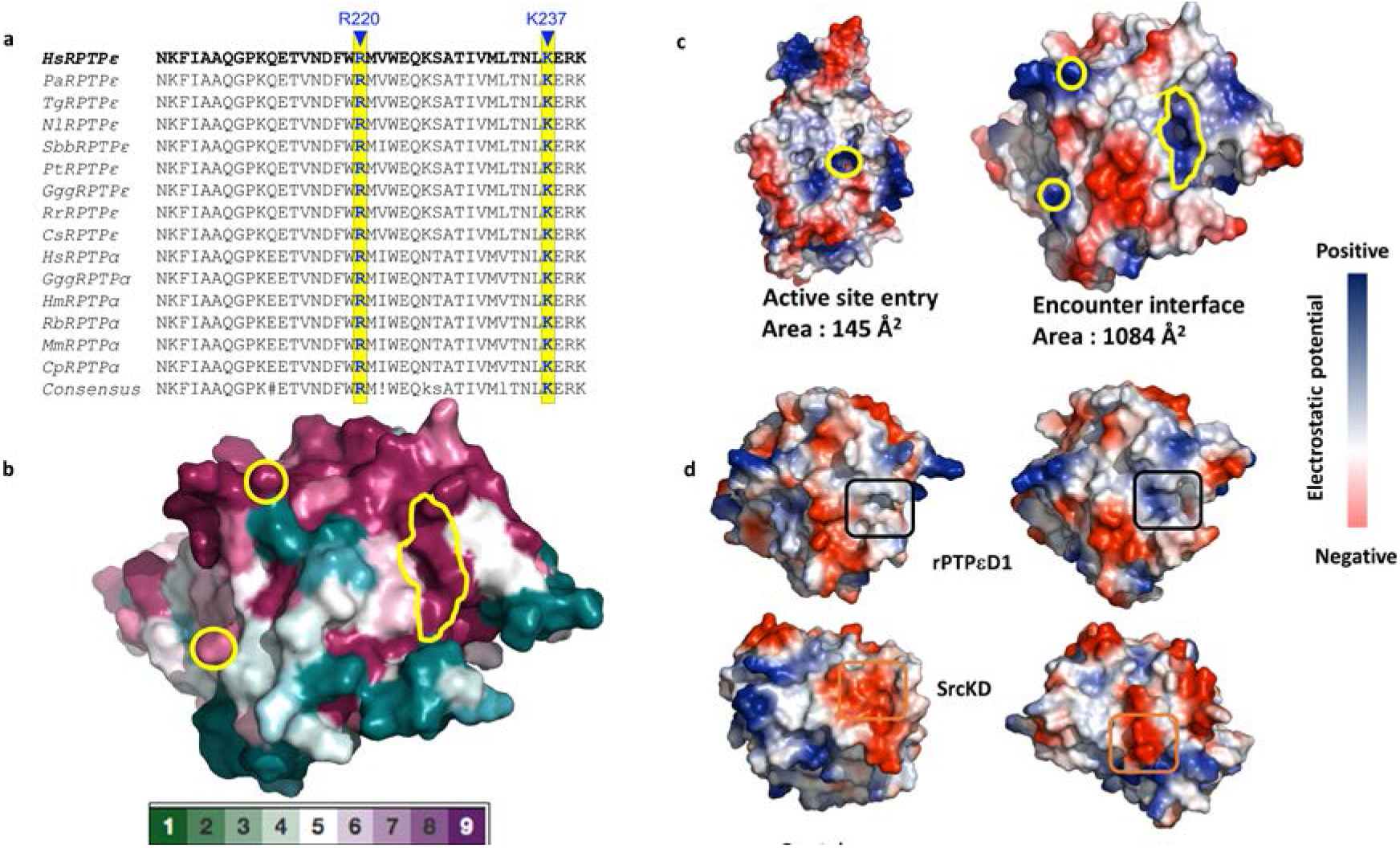
Activity comparison of rPTPεD1/rPTPεD1-R220E and surface potentials analyses. (a) Multiple sequence alignment of rPTPε and rPTPα from different species. The interface residues of the rPTPεD1: SrcKD complex, R220 and K237 are conserved in all available species of rPTPε/α. (b) The encounter residues are shown in surface representation with evolutionarily conserved and divergent residues colored. Fully conserved residues are in pink and highly divergent residues are in green. The encounter interface is highlighted in yellow circles. (c) The surface area comparison between the active site entry (left) and the encounter interface (right). The active site entry of pTyr and the encounter interface are highlighted in yellow circles. (d) Electrostatic potential surface focusing on encounter interface of rPTPεD1 and SrcKD from the initial crystal structure (left) and final MD model (right). The positive surface colored in blue and the negative surface colored in red. The rPTPεD1-R220 and SrcKD-E486 regions are highlighted in black and orange squared

### Structure analyses of the encounter interface and R220/E486 conformations

To analyze the rPTPεD1: phospho-Src complex interface, we compared an electrostatic surface potential map of the crystal structures and simulated complex. Compared to the rPTPεD1 active site whose entry covers a surface of 145 Å^2^, the encounter interface provides additional attractive interaction surface area of 1084 Å^2^ (Fig. 5c). The comparison of individual structures from the crystal and simulated complex revealed that the simulated complex undergoes local conformational changes near the encounter interface and leads to favorable electrostatic interactions (Fig. 5d and S2). In particular, the residue R220 at encounter interface adopts an alternative rotamer conformation (favored 11.3% based on Molprobity analysis), providing an attractive favorable contact in the interface. In the case of SrcKD, Glu486 locating on a flexible loop flips up to a distance of 4.5 Å towards the R220, contributing to a favorable electrostatic interaction in the encounter complex interface. Further analyses of uncomplexed ensembles from our simulated dissociate pathway, several rPTPεD1-R220 conformers similar to the crystal structure can be observed and the Src-loop containing E486 is dynamic, suggesting that the crystal structure can change freely to the conformation of simulated MD complex (Fig. S4).

## DISCUSSION

Although X-ray crystallography and cryoEM are best suited for protein complex structure determination, the weak and transient nature of interactions between PTPs and their protein substrates has substantially hindered structural understanding of these complex interactions using conventional approaches. Multiscale MD simulation provides an alternative way to reveal these type of interactions but requires extremely long time-scale simulation to reach convergence. In this study, we describe a general approach to probe the interactions using rPTPεD1: phospho-SrcKD complex as a model system. Our strategy, which employs small-angle X-ray scattering guided docking and pTyr-tailored molecular simulation, revealed to date unknown interactions distal to the rPTPεD1 active site that play critical roles in complex formation. An initial complex model can be quickly provided by a SAXS guided rigid-body approach. After manual phospho-peptide docking to link the two proteins with a strong attractive force, MD relaxation guided by energy minimization fine-tune the encounter interface. *In silico* dissociation sampling after purposely removing pTyr interaction allows identification of the key interactions which can then be validated by site-mutagenesis and developed binding experiments. Our integrative approach provides a strategy for the structural characterization of other PTP: phospho-protein complexes.

Our structure reveals a key charge-charge interaction between rPTPεD1-R220 and phospho-SrcKD-E486 far from the active site for complex formation (Fig. 3f). Systematic analyses of 131 protein-protein hetero-complexes in the PDB also shows that transient charge-charge interaction is predominant in signaling complexes, which is consistent with our finding^29^. The electrostatic interactions remain effectively with a distance of 10-20 Å^30^. We postulate a long-range electrostatic interaction between R220 and E486 brings rPTPεD1 and SrcKD into proximity at the beginning of complex formation. Once the R220:E486 encounter interface is established, the second interaction between K237 and D518 is formed. The conformation adopted by D518 in the K237: D518 interaction orients the dynamic C-terminal pTyr527 (connected through the main chain) into the rPTPεD1 active site for dephosphorylation. This proposed association pathway accompanies a 7.5-fold wider charge-charge interface to increase the probability of rPTPεD1: phospho-SrcKD complex formation compared to the interface between rPTPεD1 and pTyr (Fig. 5e). The R220E variant that partially disrupts the charge-charge interaction and additional contacts at the encounter interface results in the reduction of the association rate and phosphatase activity, supporting our proposed mechanism.

Structure analyses suggested that individual protein conformations in the un-complexed state (crystal structure) and the complexed state (MD simulated ones) can be freely interchangeable prior to complex formation. However, conformations in the complexed state provide more favorable binding energies by increasing attractive interactions. In contrast, the R220E variant mimicking the repulsive interaction that would result from the rigid-body docking conformation observed in the crystal structure binds to Src with a slower association rate and has reduced phosphatase activity. The R220E variant and wild-type have similar *k*_off_ rates. Taken together, our data demonstrate that the molecular recognition of rPTPεD1 is consistent with a conformational selection mechanism (Fig. 6), with a conformational change before the binding event, rather than an induced-fit mechanism.

**Figure 6.**
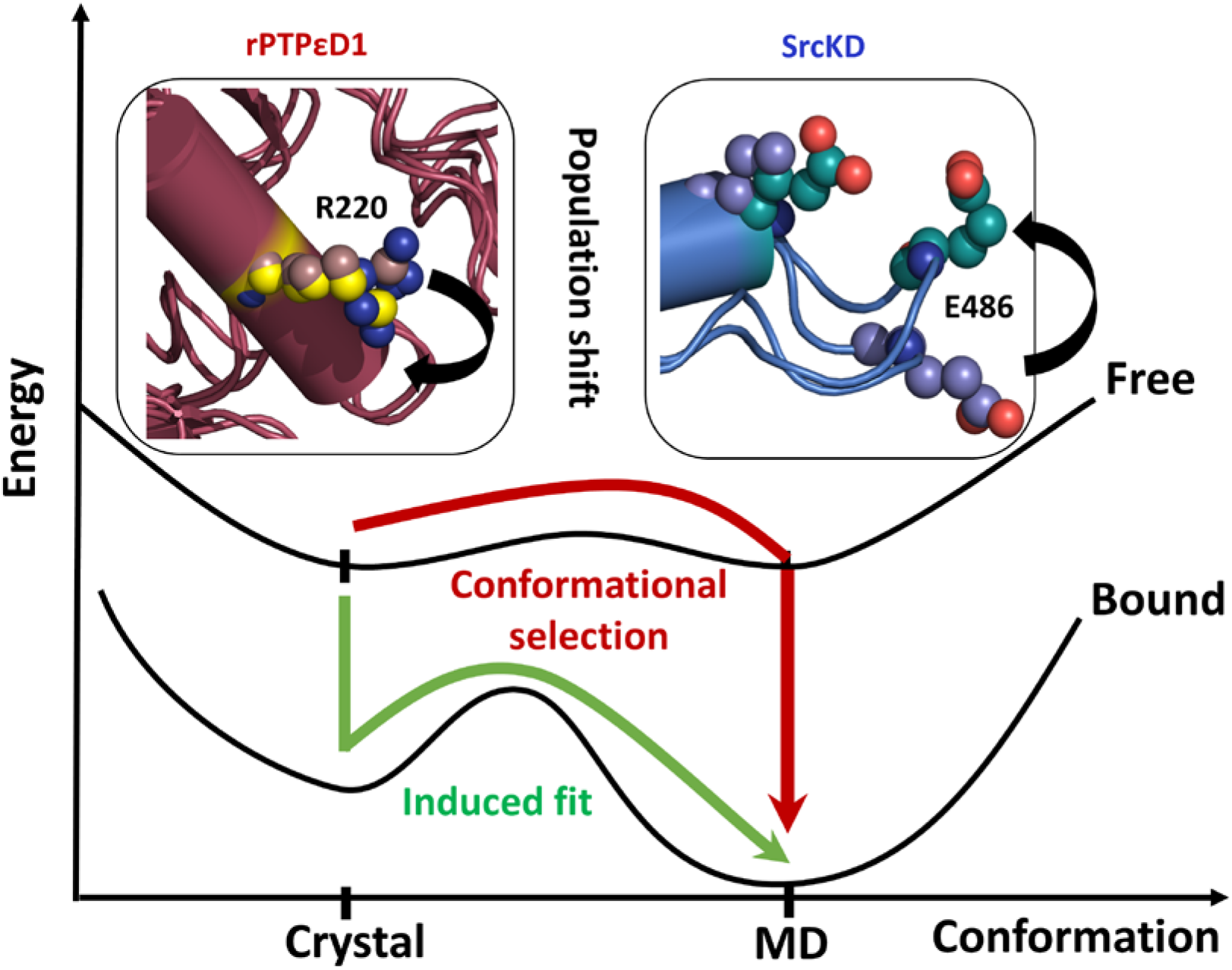
Proposed conformational selection mechanism. The schematic diagram illustrates the conformational selection of the rPTPεD1: Phospho Src complex.

## Materials and Methods

### Cloning, expression and purification of the rPTPεD1 and SrcKD

His tagged human Src kinase domain (Trp 260-Leu 533) was cloned into a modified pET vector for recombinant protein production. In this construct, the ATP binding site (K295M) and autophosphorylation sites (Y416F) of SrcKD were mutated to generate fully inactive Src by site-directed mutagenesis. The His- and MBP-tagged human rPTPεD1 Domain (Ser 101-Thr 400) was cloned into a modified pET9a vector. A TEV protease cutting site is located between the tag and constructed protein domains. In this study, the substrate trapping C335S mutant was created for all binding studies. The R220E mutant was generated by site-directed mutagenesis. The human rPTPαD1 (Ser211-Thr512) plasmids were created in a similar way to the rPTPεD1 constructs. The plasmids were subsequently transformed into *Escherichia coli* BL21 (DE3) cells, which were grown at 37 °C in LB medium supplemented with 100 mg/L ampicillin until OD_600_ reached ~ 0.8. The proteins were overexpressed by the addition of IPTG to a final concentration of 0.5 mM. After an additional incubation for 8 hours at 16 °C, the culture was harvested by centrifugation at 8,000 × g for 20 minutes. Cell pellets were re-suspended in buffer-A (100 mM phosphate buffer at pH 7.5, 500 mM NaCl, 10% glycerol and 10 mM β-mercaptoethanol). The suspension was lysed by sonication and centrifuged at 35,000 × g at 4 °C for 45 minutes. The supernatant was loaded onto a nickel-affinity column pre-equilibrated with buffer-A. The protein was washed with buffer-A with various concentrations of imidazole and finally eluted with buffer-B (50 mM Tris-HCl pH 7.5, 500 mM NaCl, 500 mM imidazole). The fractions containing tagged SrcKD or rPTPεD1 were pooled and treated with TEV protease to remove the tags. The tag-free SrcKD/rPTPεD1 proteins were further purified by a nickel-affinity column. The flow-through was subsequently concentrated and purified by size-exclusion chromatography (SEC) column, which was pre-equilibrated with 20 mM Tris pH 7.5, 150 mM NaCl, 5 mM DTT, 5% glycerol.

### The Phosphorylated rPTPεD1: SrcKD Complex

The phosphorylation of 60 μM SrcKD kinase was achieved by His- and chitin-binding domain (CBD)-tagged Csk in the presence of 10 mM MgCl2, 10 mM DTT, 45 μM ATP, and 5% glycerol at room temperature for 30 minutes and was separated from His-CBD-CSK protein using Chitin beads. The phosphorylated SrcKD was successively tested for the extent of phosphorylation by ^32^[P] assay^31^. The phosphorylated SrcKD was mixed with 0.5 molar excess of rPTPεD1 and the resultant complex was loaded onto a Superdex 200 HR 10/300 Increase column and separated using 20 mM Tris pH 7.5, 150 mM NaCl, 5 mM DTT buffer. The formation of the rPTPεD1: SrcKD complex was further confirmed not only be elution volume but also SDS-PAGE electrophoresis. Similarly, a comparison of rPTPεD1/rPTPεD1-R220E towards cSrc was performed using a Superdex 75 HR 10/300 column.

### rPTPεD1-Src-peptide complex

The FiTC labeled pTry527 phosphorylated Src-peptide (EAQpYQPGENL) was synthesized at the in-house peptide-synthesis facility. The peptide was dissolved in DMSO to a final concentration of 10 mM. The rPTPεD1 (50 μM) was incubated with pTyr527 phosphorylated Src-peptide (150 μM) at room temperature for 1 hour. The subsequent complex was separated using a Superdex 200 HR 10/300 (Increase) column, pre-equilibrated with SEC buffer. The absorbance was monitored at 280 nm and 494 nm for protein and FiTC detection, respectively.

### Analytic ultracentrifugation

Sedimentation velocity analysis was performed with an XL-A analytical ultracentrifuge (Beckman Coulter) with absorption optics, using an AnTi60 rotor. Samples in 20 mM Tris-HCl at pH 7.5, 50 mM NaCl and 1 mM TCEP were added to double-sector centerpieces and centrifuged at 45,000 rpm for 18 hours at 20 °C or 4 °C. Detection of concentrations as a function of radial position and time was performed by optical density measurements at a wavelength of 280 nm, with absorbance profiles recorded every 3 min. The buffer density and viscosity were calculated using the software SEDNTERP, and the data were analyzed using SEDFIT software. The plots of the AUC profiles were made with Prism7.

### SAXS model of rPTPεD1: phospho-SrcKD Complex

Small-angle X-ray scattering (SAXS) data was collected on beamline BL23A at the National Synchrotron Radiation Research Center (NSRRC, Hsinchu, Taiwan). The rPTPεD1: phospho-SrcKD complex was prepared in 20 mM Tris pH-8.0, 50 mM NaCl, 5 mM DTT and concentrated to 6 mg/ml, then was injected to online SEC-SAXS equipped with a temperature-controlled (15 °C) silica-based SEC column (Agilent BioSEC-3)^32^. The SAXS profiles of sample buffer after the elution peaks of protein samples were collected for background subtraction. All SAXS two-dimensional images were processed and transferred into one-dimensional intensity curves by an in-house program established by LabVIEW^32^. The output text files were further processed using the PRIMUS software suite^33^. Parameters such as radius of gyration (*R*_g_), the maximum particle dimensions (*D*_max_), and the Porod volume (*V*_p_) were evaluated using standard procedures (Table S1)^33^. The program GNOM was used to calculate the distance distribution function. The reported crystal structures of the SrcKD, rPTPεD1 were used as a template to dock the complex structure using the CORAL program^20^. The pTyr527 was manually docked to rPTPεD1 active site based on the crystal structure of pTyr-peptide bound PTP1B using COOT^25,34^. The missing residues were added using RosettaCommons^35^. By holding the rPTPεD1 and pTyr527 region of SrcKD connected and allow the missing N-terminus residues to be flexible, the docking model was improved by conformational sampling followed by SAXS validation using BIBLOMD^36^. The quality of the fit between models and experimental SAXS profile was calculated by FoXS ^37^.

### Encounter Interface optimization by Molecular Dynamics Simulation

To understand the key residues involved in the encounter interface, molecular dynamics (MD) simulations were conducted using the Amber 16 package^38^. The rigid-body model of rPTPεD1: phospho-SrcKD complex was taken as a starting coordinate for MD simulations. MD Simulations were performed based on a force field Amber ff14SB ^39^ that extends the improved residue side-chain torsion potentials. The residue Mulliken charges were calculated based on the libraries in the Amber 16 package. Periodic boundary conditions were imposed with box lengths of 98.33 × 151.56 × 105.79 Å^3^, containing 592 amino acid and 48655 TIP4P water models. The MD System underwent a 15 ns annealing process under the constant pressure of 1.0 bar with equilibrated steps from 0 to 300 K. The constrain force applied to residue pairs varied from 200 kcal/mol to 5 kcal/mol until the system density was ~ 1.0 ± 0.01 g/cm^3^. A Langevin thermostat was used to maintain the system temperature by controlling the collision frequency at 1 ps^−1^ to the target temperature 300 K. MD simulations were carried out in the canonical ensemble (NVT) with the Langevin thermostat to maintain the system temperature. The SHAKE algorithm was implemented to constrain the covalent bond including hydrogen atoms. Numerical integration was performed with a time-step of 1 fs for all MD simulations. We performed ∼0.3 μs MD simulations for checking systems equilibrium and analyze.

### Searching the Pathway Trajectory with Steered MD Simulations

We performed steered MD (SMD) simulations^40–42^ for the dissociated and associated state of the complex, employing distance-based collective variables between the rPTPεD1 and SrcKD domain with the center of mass (COM) distance ~ 44.5 – 62.5 Å. The force weight sets on the x, y and z-component with the restrain force equal to 5 kcal mol^−1^ Å^−2^. For the dissociation process, the initial model was taken from the MD optimized structure. Simulations were carried out for every 36 windows of 1 ns run, via a strain velocity during sampling relaxation (pulling force: 5 kcal mol^−1^ Å^−2^, velocity: 0.0005 Å ps^−1^), corresponding to a total simulation time of 0.36 μs. For the association process, the started model was taken from the trajectory of the dissociation process, with a COM distance ~57 Å. Simulations were carried out for every 50 windows of 1 ns run, via a slower strain velocity during sampling relaxation than that of the dissociation process (pulling force: 5 kcal mol^−1^ Å^−2^, velocity: 0.00025 Å ps^−1^), corresponding to a total simulation time of 0.5 μs.

### Estimating the Free Energy via Potential Mean Force (PMF) and Free Energy Pathway (FEP)

In umbrella sampling (US-PMF)^43–45^, harmonic restraint is placed at successive points along with the reaction coordinate with restraining potential form V(t) = k(x_t_-x_o_)^2^, where x_0_ is the target distance and k is the force constant. The reaction path was stratified into a series of intermediate windows, ranging from 44.5 to 62.5 Å for the separation of rPTPεD1 and SrcKD. Instantaneous values of the force were accrued in bins of width equal to 0.5 Å for separation PMFs. The reaction coordinate is defined as the z-component of the center-of-mass (COM) distance between the rPTPεD1 and the SrcKD. The path is divided into 36 windows at ~0.5 Å intervals. The restraint force constants: 5 kcal/mol Å^2^ to ensure overlap between each rPTPεD1/SrcKD window and each window is simulated for 1 ns, corresponding to a total simulation time of 0.36 μs. After the simulation, the free-energy curves are combined by WHAM which is used to convert the probabilities into the PMF along with the reaction coordinate at 300 K. The number of points in the final PMF was 1800 (0.01 Å for 1 bin) and the convergence tolerance for the WHAM calculations was 0.001 kcal/mol.

To decompose the estimated binding free energy from US simulations, two-end-state free energy calculations were performed directly based on the trajectories derived from the US simulations. We employed the MMGB/PBSA^46,47^ method to decompose the binding free enthalpy. The electrostatic solvation energy was calculated by the GB model developed by Onufriev (igb=2)^46^. The exterior dielectric constant was set to 80 and the solute dielectric constant was set to 0.1. The non-polar contribution of the desolvation energy (DGSA) was estimated from the solvent accessible surface area (SASA) using the LCPO algorithm.

### BLI binding assay

To perform the binding experiments, the rPTPεD1-C335S, rPTPεD1-C335S/R220E and phospho His-cSrcKD-K295M/Y416F proteins were purified. The binding kinetics of rPTPεD1 and phospho cSrcKD association was measured by BLItz using Ni-NTA biosensor tips (ForteBio Inc.). Further, the Ni-NTA sensors were pre-hydrated for 10 minutes in SEC buffer. The bait, phospho-His-SrcKD at a concentration of 58 µM was immobilized to Ni-NTA sensor tips for 3 min. To maintain stable phosphorylation of SrcKD, the buffer was supplemented with fresh 100 μM ATP and 1 μM Csk before every immobilization. Once the bait protein reached saturation, subsequent association of rPTPεD1/rPTPεD1-R220E proteins to the bait were allowed for 120 sec followed by a 3 min. dissociation step. The BLI data was processed by the BLItz Pro software and plotted in Graphpad Prism7.

### pNPP assay

The rPTPεD1/rPTPαD1 phosphatase assay using pNPP as the substrate was performed as previously described^48,49^. In brief, the purified rPTPεD1, rPTPαD1, rPTPεD1-R220E, or rPTPαD-R317E proteins were added in a solution containing 20 mM Tris-HCl (pH 7.5), 50 mM NaCl, and 20 mM pNPP. The reactions were incubated for 30 min, the level of dephosphorylation was measured at 405 nm using a UV spectrometer. All measurements were performed in triplicate.

### Phosphatase activity assay

The phosphatase activity of rPTPεD1/rPTPαD1 wild type and rPTPεD1-R220E/rPTPαD1-R317E mutants were measured using the phosphate colorimetric assay kit (Bio Vision, Milpitas, CA). The purified wild-type rPTPεD1/rPTPαD1 and rPTPεD1-R220E/rPTPαD1-R317E were mixed with 20 μM phosphorylated cSrcKD in a solution containing 20 mM Tris-HCl (pH 7.5), 50 mM NaCl and incubated for 30 min. The dephosphorylation of the phosphatase was quenched by the addition of kit reagent, the reactions were further incubated for 15 min. The dephosphorylation levels were measured by absorbance at 650 nm using a TECAN M1000pro. Assays were performed according to the manufacturer’s instructions. All measurements were performed in triplicate. A control phosphorylated cSrcKD was subtracted from all runs. The figures for publication were generated using Graphpad Prism7 software.

## Data availability

The SAXS data accession code for rPTPεD1: phospho-SrcKD complex is SASDJ33.

## Acknowledgment

We thank Dr. Orion Shih and Kuei-Fei Liao at the NSRRC TLS23A beamline and Dr. Meng-Ru Ho of the Biophysics Facility, Institute of Biological Chemistry, Academia Sinica for assistance in different data collection. We acknowledge the peptide synthesis core, Institute of Biological Chemistry, Academia Sinica for pTyr-peptide synthesis and the National Synchrotron Radiation Research Center, Taiwan, for use of the BL23A beamline for SEC-SAXS data collection. We also thank Dr. Yu-Ching Lin for his assistance in BLI data interpretation. This work was partially funded by the Academia Sinica (AS-104-TP-B05) and Taiwan Protein Project (Grant No. AS-KPQ-105-TPP).

## Author Contributions

N.E.K and S.K.T performed recombinant protein preparation. N.E.K and Y.Y collected SAXS data and completed the complex model building. C.H.Y. performed MD simulations. N.E.K performed phosphatase activity and binding assays. N.E.K., C.H.Y., H.C.Y and M.C.H. wrote the manuscript. N.E.K., H.C.Y and M.C.H. designed the experiments. H.C.Y and M.C.H. supervised the work.

## Supplementary figures and tables

**Figure S1.**
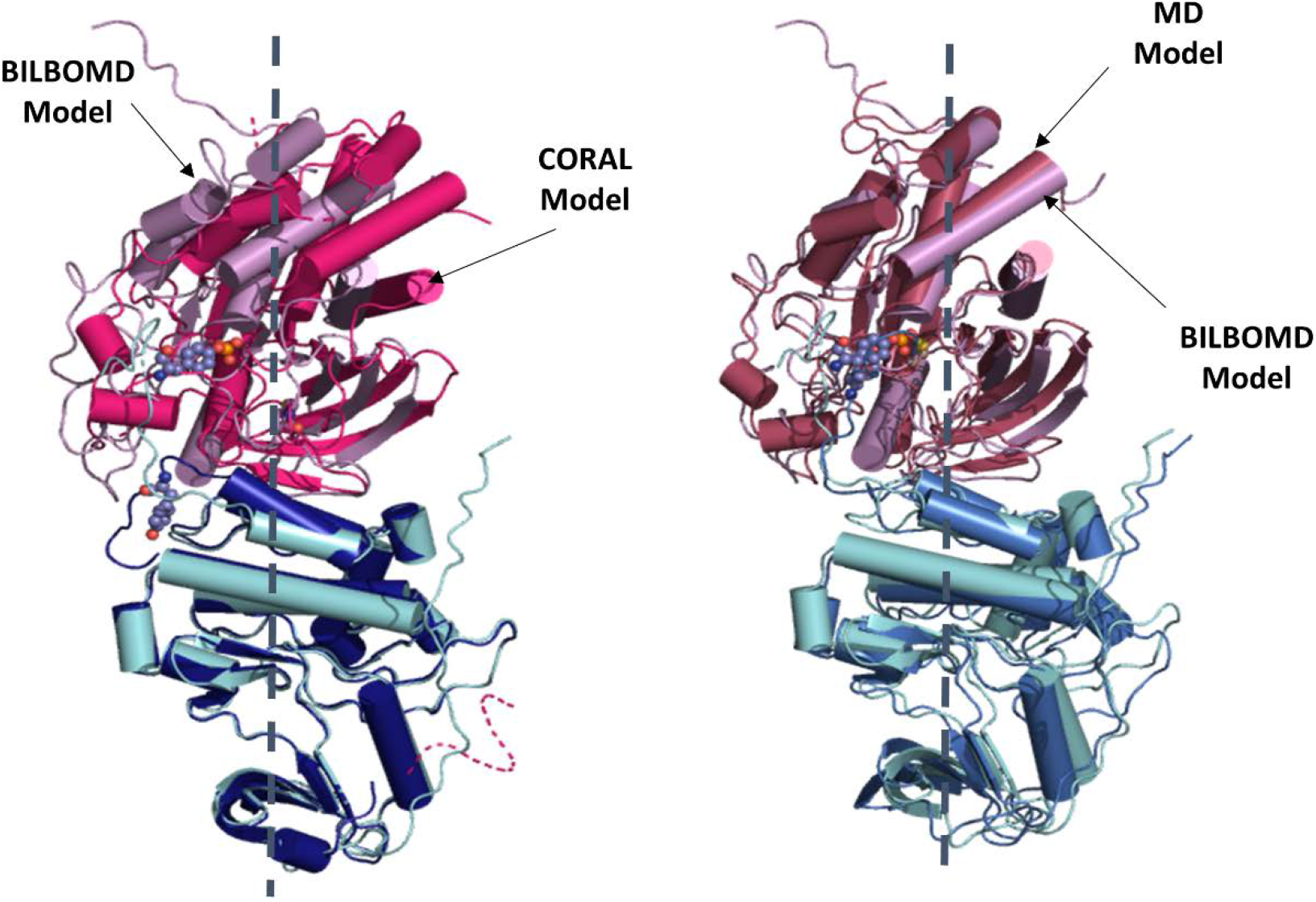
Comparison of CORAL, BILBOMD and MD simulation model of rPTPεD1: phospho-SrcKD complex. The overlaid structures of CORAL and BILBOMD are shown on the left. The overlaid structures of BILBOMD and MD model are shown on the right.

**Figure S2.**
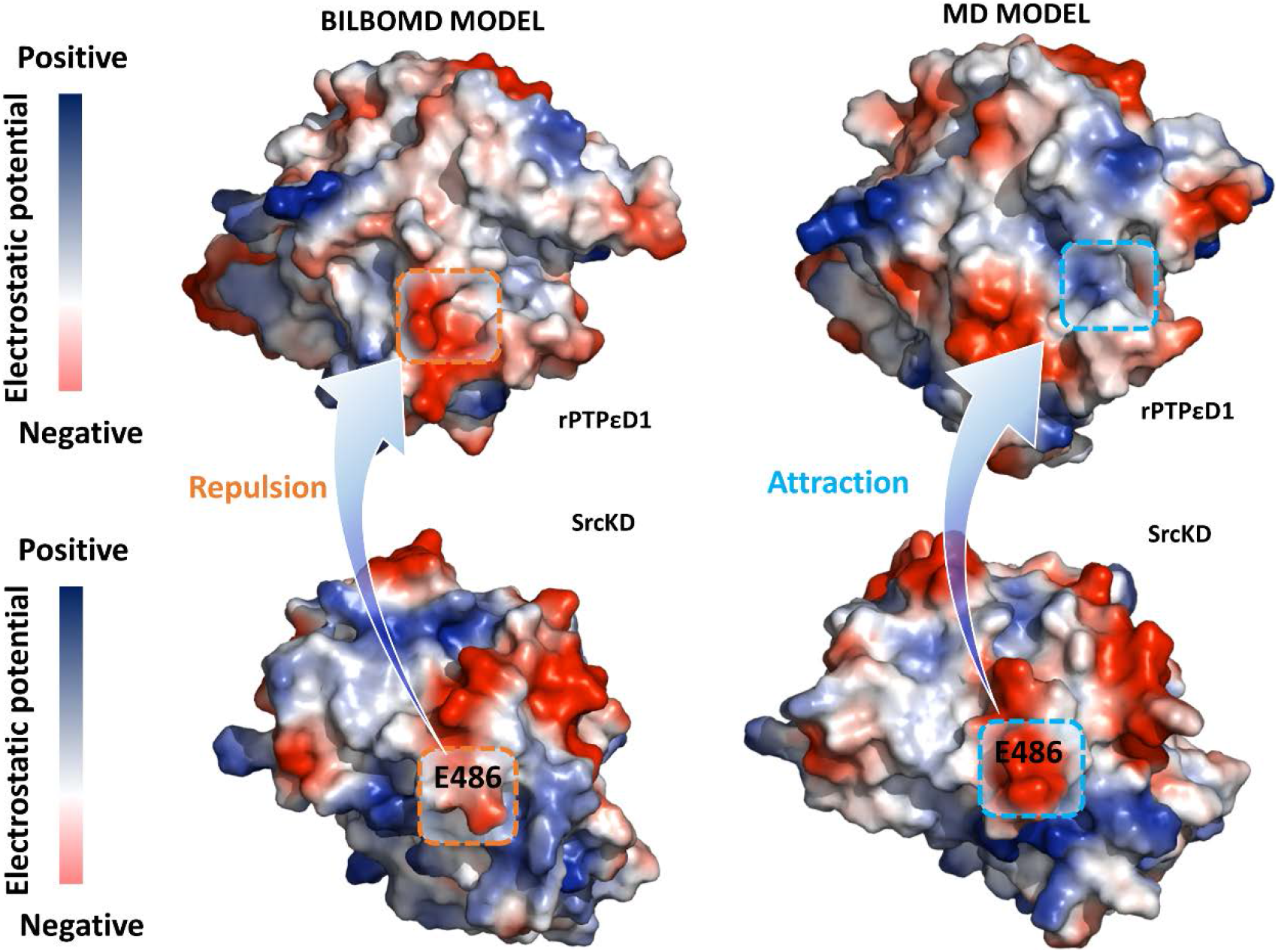
The comparison of electrostatic potential surface focusing on encounter interface of rPTPεD1 and SrcKD between BILBOMD and final MD model. The positive surface colored in blue and the negative surface colored in red. The repulsion region (orange square) and attractive region (blue square) show the interaction of the encounter surface between rPTPεD1 and SrcKD.

**Figure S3.**
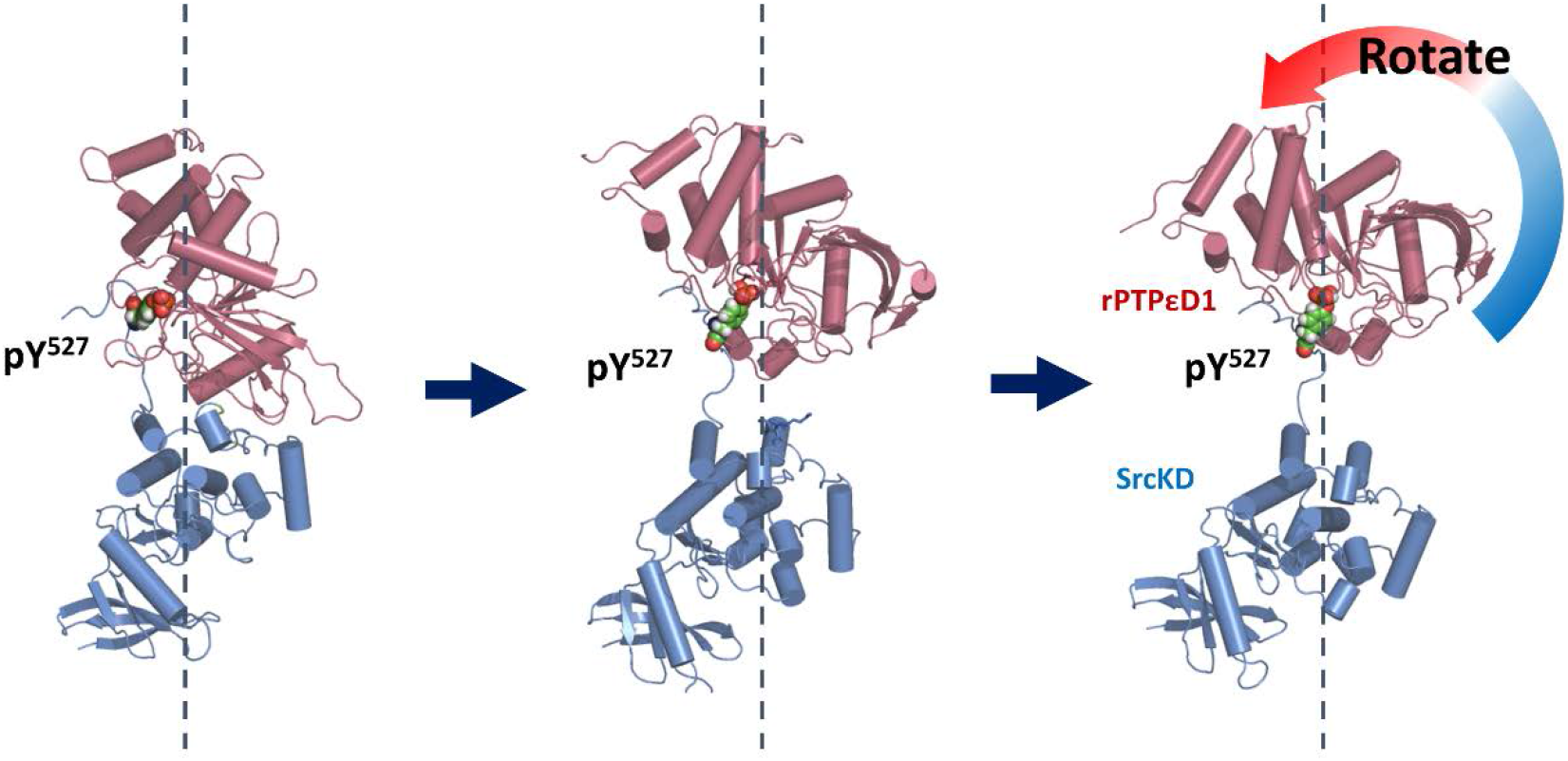
Illustration of how a strong interaction between pTyr 527 and rPTPεD1 causes unreasonable dissociation sampling. During dissociation trajectory, pTyr527 of SrcKD remains bound to the rPTPεD1 active site, causing the rotation of rPTPεD1.

**Figure S4.**
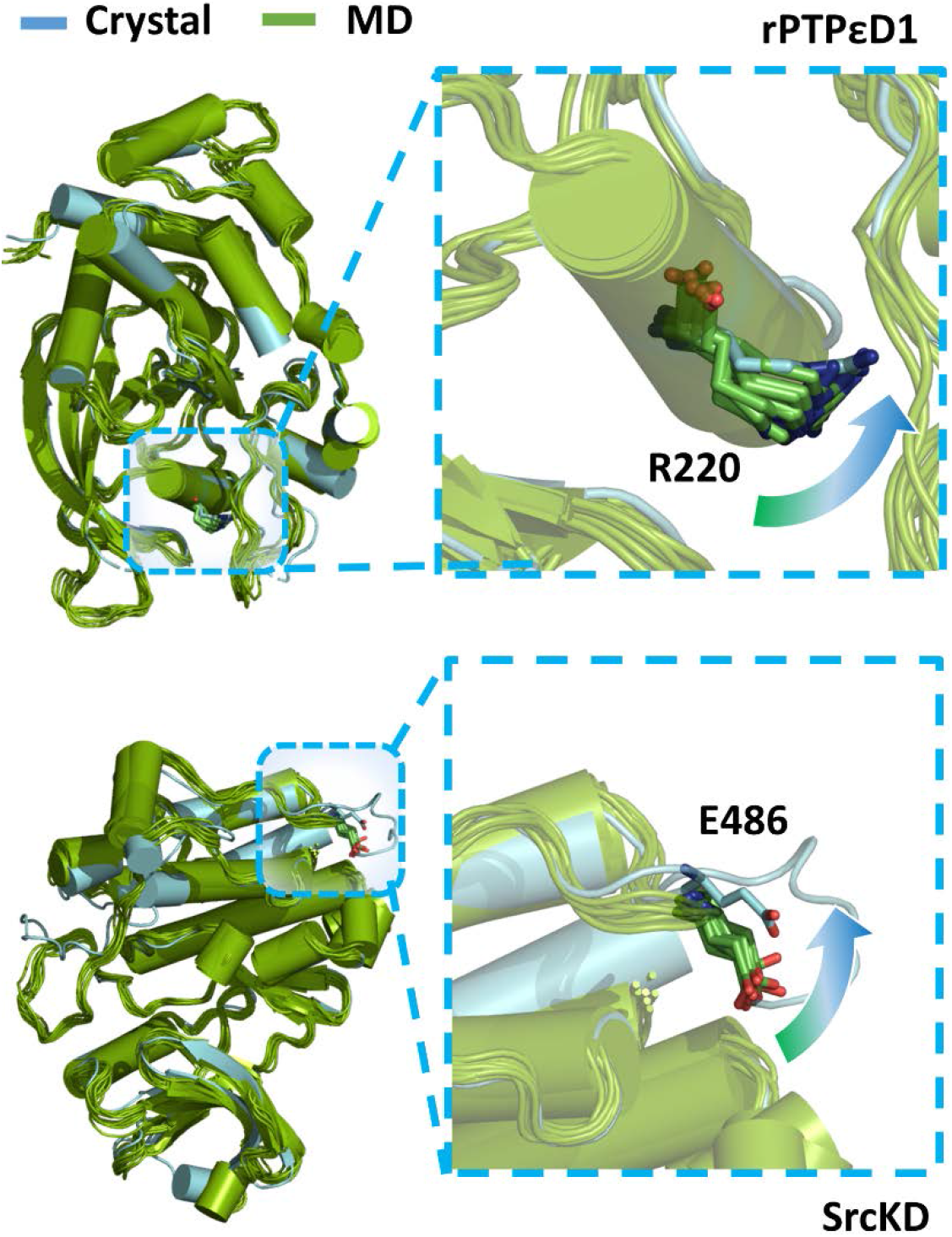
The schematic diagram displays the MD simulation trajectories of the unbound form of rPTPεD1 and Src.

**Table S1.**
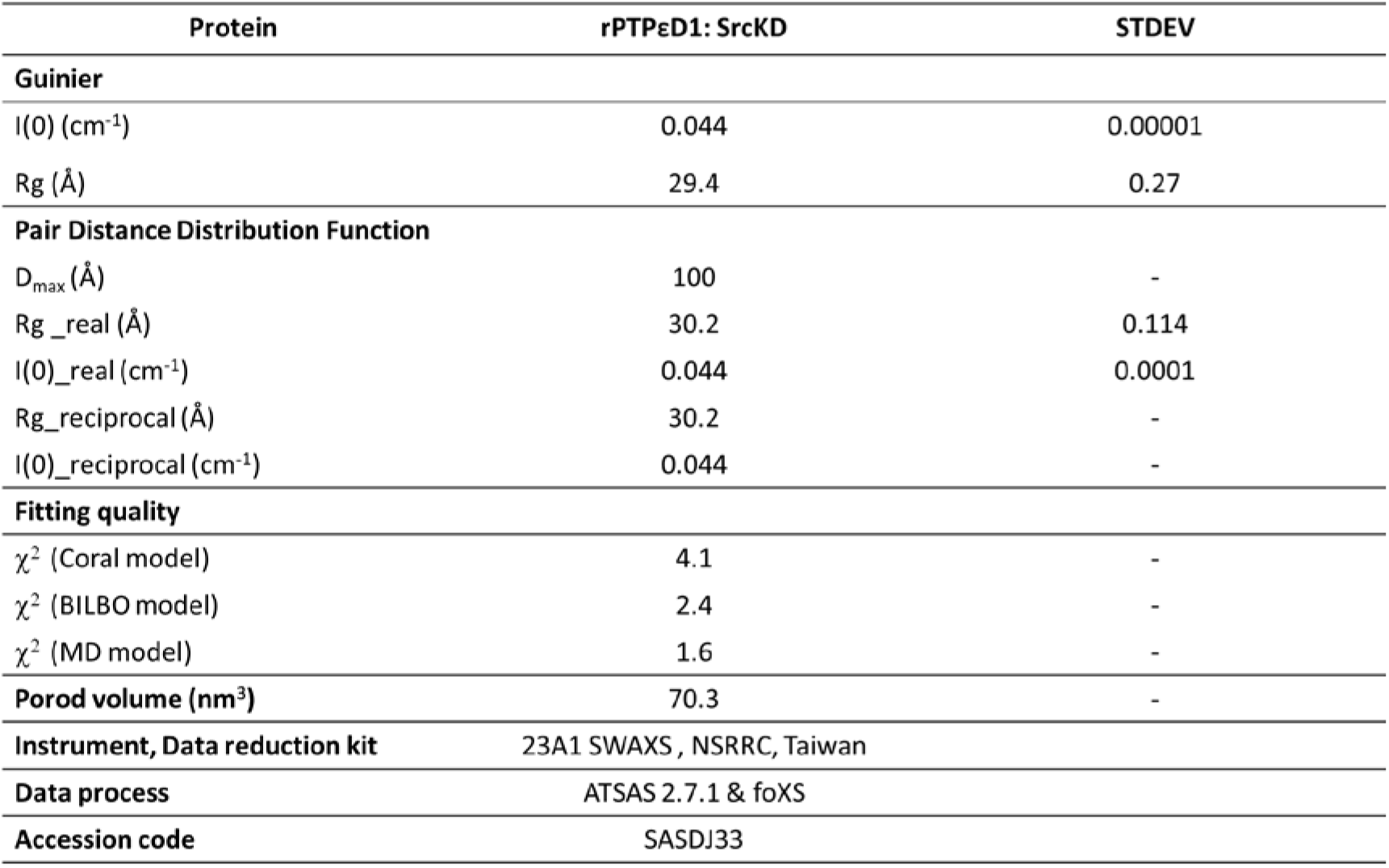
SAXS data table and analysis parameters for rPTPεD1: cSrcKD complex.

| Protein | rPTPεD1: SrcKD | STDEV |
| --- | --- | --- |
| **Guinier** |  |  |
| I(0) (cm<sup>-1</sup>) | 0.044 | 0.00001 |
| Rg (Å) | 29.4 | 0.27 |
| **Pair Distance Distribution Function** |  |  |
| D<sub>max</sub> (Å) | 100 | - |
| Rg_real (Å) | 30.2 | 0.114 |
| I(0)_real (cm<sup>-1</sup>) | 0.044 | 0.0001 |
| Rg_reciprocal (Å) | 30.2 | - |
| I(0)_reciprocal (cm<sup>-1</sup>) | 0.044 | - |
| **Fitting quality** |  |  |
| χ<sup>2</sup> (Coral model) | 4.1 | - |
| χ<sup>2</sup> (BILBO model) | 2.4 | - |
| χ<sup>2</sup> (MD model) | 1.6 | - |
| **Porod volume (nm<sup>3</sup>)** | 70.3 | - |
| **Instrument, Data reduction kit** | 23A1 SWAXS, NSRRC, Taiwan |  |
| **Data process** | ATSAS 2.7.1 & foXS |  |
| **Accession code** | SASDJ33 |  |

**Table S2.** Comparison of the form factor characteristics of rPTPεD1, SrcKD and complex in the MD-derived model and rigid-body model.

|  | MD-derived |  |  | BILBOMD |  |  | Experimental |
| --- | --- | --- | --- | --- | --- | --- | --- |
|  | rPTPε | SrcKD | Complex | rPTPε | SrcKD | Complex | Complex |
| **Rg (Å)** | 19.9 | 22.4 | 30.5 | 20.0 | 22.4 | 30.3 | 29.4 |
| **Minor (Å)** | 17.3 | 19.6 | 21.2 | 17.9 | 21.1 | 20.5 | 20.7 |
| **Major (Å)** | 15.6 | 19.6 | 23.3 | 16.1 | 19.0 | 22.6 | 22.8 |
| **n/m<sup>a</sup>** | 0.9 | 1.0 | 1.1 | 0.9 | 0.9 | 1.1 | 1.1 |
| **Length (Å)** | 56.5 | 64.2 | 91.3 | 56.9 | 64.9 | 91.1 | 90.7 |

<sup>a</sup>n/m is the major/minor axis of the elliptical cylinder model.

